# Microglial MyD88-dependent pathways are regulated in a sex-specific manner in the context of HMGB1-induced anxiety

**DOI:** 10.1101/2024.04.22.590482

**Authors:** Ashleigh Rawls, Dang Nyugen, Julia Dziabis, Dilara Anbarci, Madeline Clark, Kafui Dzirasa, Staci D. Bilbo

## Abstract

Chronic stress is a significant risk factor for the development and recurrence of anxiety disorders. Chronic stress impacts the immune system, causing microglial functional alterations in the medial prefrontal cortex (mPFC), a brain region involved in the pathogenesis of anxiety. High mobility group box 1 protein (HMGB1) is an established modulator of neuronal firing and a potent pro-inflammatory stimulus released from neuronal and non-neuronal cells following stress. HMGB1, in the context of stress, acts as a danger-associated molecular pattern (DAMP), instigating robust proinflammatory responses throughout the brain, so much so that localized drug delivery of HMGB1 alters behavior in the absence of any other forms of stress, i.e., social isolation, or behavioral stress models. Few studies have investigated the molecular mechanisms that underlie HMGB1-associated behavioral effects in a cell-specific manner. The aim of this study is to investigate cellular and molecular mechanisms underlying HMGB1-induced behavioral dysfunction with regard to cell-type specificity and potential sex differences. Here, we report that both male and female mice exhibited anxiety-like behavior following increased HMGB1 in the mPFC as well as changes in microglial morphology. Interestingly, our results demonstrate that HMGB1-induced anxiety may be mediated by distinct microglial MyD88-dependent mechanisms in females compared to males. This study supports the hypothesis that MyD88 signaling in microglia may be a crucial mediator of the stress response in adult female mice.

## Introduction

Anxiety is a common cause of mental health-related disability. Anxiety disorders describe a class of disorders in which the affected person experiences consistent and often unfocused anxiety and worry (Bandelow and Michaelis., 2015). Though generally high in prevalence in both sexes, anxiety is twice as common in women throughout their lifetime (Mclean et al., 2011). Few biological mechanisms have been elucidated that explain the increased prevalence of depression or anxiety in women; however, we do understand that stress is a pivotal factor in the development and recurrence of anxiety disorders regardless of sex or age (McEwen and Akil, 2020).

Stress instigates many alterations throughout the brain-body axis (Fonken et al., 2018). Specifically, changes in the peripheral and central immune systems have been observed in patients with anxiety and comorbid depression (Costi et al., 2021). Preclinical models also show immune responses to physical and psychological stressors (Ménard et al., 2017). Psychological stress acts as a sterile immune challenge that elicits proinflammatory responses centrally and peripherally (Hodes et al., 2021). Sterile immune responses increase the release of danger-associated molecular patterns (DAMPs), which are recognized by conserved pattern recognition receptors (PRRs) primarily on immune cells. This response has been demonstrated to be an essential mechanism by which stress elicits behavioral changes (Weber et al., 2018). High mobility group box one (HMGB1) is one such DAMP released during stress that activates multiple PRRs, including toll-like receptors (TLRs) and the receptor for advanced glycation end products (RAGE) (Kang, 2014). Previous results demonstrate that stress is associated with increased HMGB1 release in the brain and subsequent activation of TLR2, TLR4, and RAGE signaling (Pluma-Pluma et al., 2023) (Chen, 2022). However, the mechanisms by which HMGB1 can impact behavioral sequelae such as anxiety in response to stressors, and thus viable therapeutic targets, remains largely undefined.

This is due in large part because nearly all cell types express HMGB1 or one of its putative receptors within the brain and little work has been done to investigate the cell type specificity of HMGB1 signaling in stress responses in the brain. Microglia, the resident innate immune cells of the brain, play key roles in proinflammatory signaling in response to stress and express PRRs at high levels relative to other CNS cells. Microglia can respond broadly to PRR activation via multiple functional alterations, including changes in phagocytic activity and cytokine release and direct interactions with neurons (Fleshner et al., 2017). Genetic deletion of TLR2/4 in microglia yields a highly stress-resilient phenotype (Nie et al., 2018). Therefore, we hypothesized that microglia have a key role in the mechanisms by which increased HMGB1 causes immune activation in the CNS and alters behavior.

To test this hypothesis, we utilized disulfide HMGB1 (dsHMGB1) locally infused into the medial prefrontal cortex (mPFC), a key stress-responsive region (Carvahlo-Netton et al., 2011) (Hinwood et al., 2012; Tynan et al., 2010). dsHMGB1 is an established pharmacological stressor that alters depressive-like behavior when circulated throughout the whole brain via direct infusion (Lian et al., 2017). Here we utilize localized administration of dsHMGB1 to incorporate a region-specific understanding of behavior with immune signaling in the context of stress. We found that dsHMGB1 locally administered to the mPFC produces a robust and reproducible anxiety-like phenotype in both male and female adult mice. We also see striking sex-specific differences in both microglial morphology as well as transcriptional activity following dsHMGB1 infusion. Finally, we found that in females specifically, conditional knockout of myeloid differentiation adaptor protein 88 (MyD88) in microglia, a key adaptor protein for many PRRs, prevents anxiety like behavior following dsHMGB1 infusion.

## Results

### HMGB1 infusion into the mPFC increases anxiety-like behavior in males and females

HMGB1 has been previously shown to increase anhedonia-like behavior (Wang et al., 2018). However, no studies to date have utilized HMGB1 as a pharmacological stressor in female subjects, calling into question how relevant sterile inflammatory mechanisms are to the stress response in females. Here, we sought first to define the impact of sub-chronic exposure to a stress-related danger signal on behavior in male and female mice. To do so, mice were infused via cannula with dsHMGB1 once daily for 5 consecutive days. Behaviors related to anxiety, sociability, and anhedonia were assessed following HMGB1 infusion into the mPFC. Anxiety-like behavior was assessed by measuring center exploration time in the open field test **(Suppl Fig 1A),** open arm exploration and latency to explore open arms of the elevated plus maze (EPM) **(Fig 1B-C)**, and latency to feed in a novelty suppressed feeding assay **(Fig 1E)**. We found that exploration of the open arms in the EPM was significantly decreased in mice treated with dsHMGB1. No changes in social preference or behavior in the tail suspension test were found **(Suppl Fig 1B-C)**. The results show that dsHMGB1 infusion to the mPFC selectively alters anxiety, and this phenotype is shared between males and females. Furthermore, we found that females specifically had a significant decrease in weight following dsHMGB1 treatment **(Fig 1D)**. We monitored estrus cycle in females on each day behavior was tested. We found no significant correlation between estrus cycle and either weight loss or novelty-suppressed feeding in response to dsHMGB1 **(Suppl Fig 2A-B)**. All remaining behavioral measures were insufficiently powered to assess impacts of estrus, due to the majority of female mice in the same part of the cycle at the time of testing. Taken together, these results demonstrate that dsHMGB1 elicits a shared behavioral phenotype of increased anxiety-like behavior in male and female mice.

**Fig. 1.**
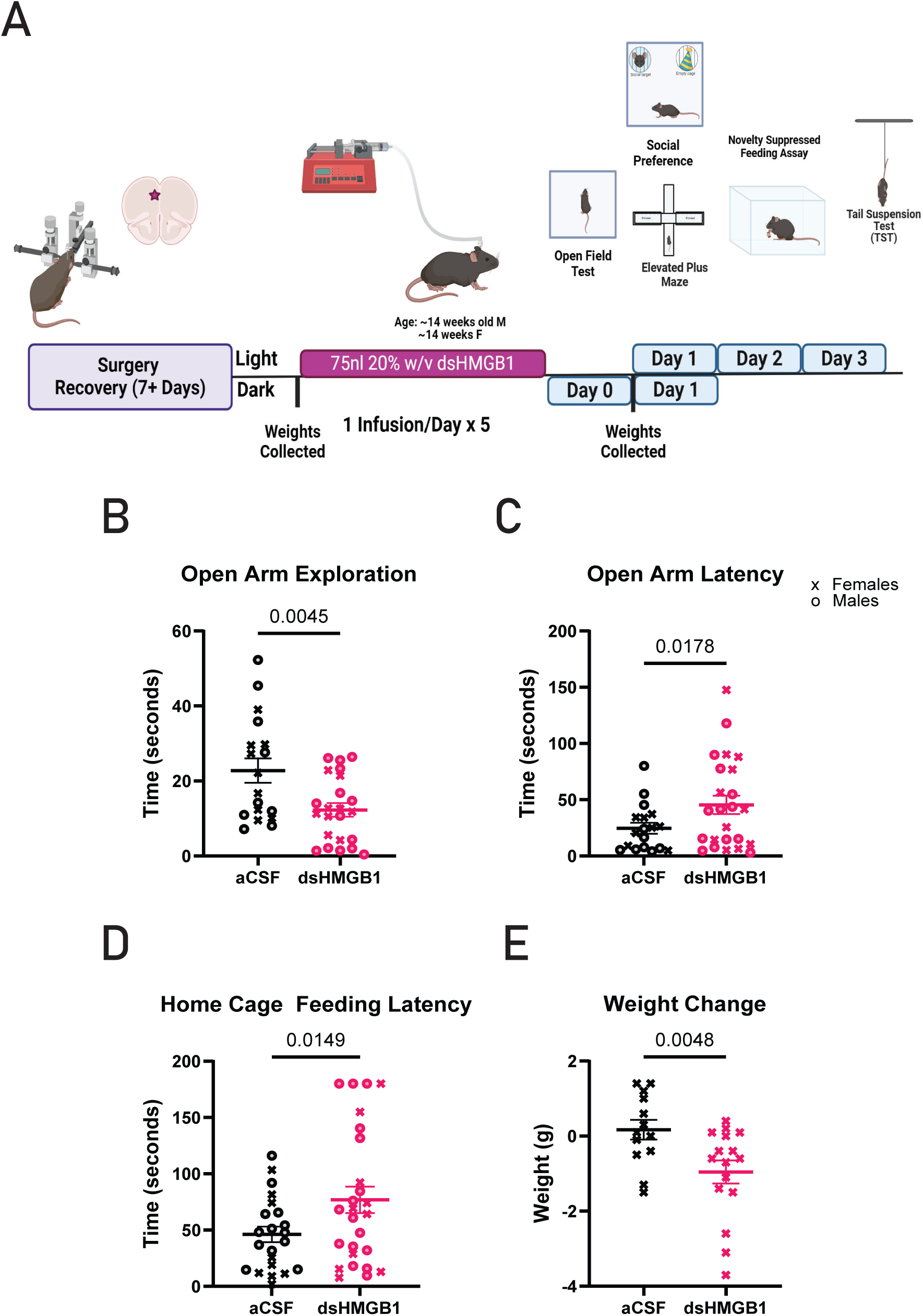
HMGB1 infusion to mPFC causes specific anxiety-like behavior. A) Schematic of Experimental Workflow (Created with Biorender) B) Open Arm Exploration C) Latency to Explore Open Arms D) Home cage Feeding Latency E) Females Show Change in Weight following Infusion Statistical analyses were performed using 2-way ANOVA to confirm primary effect of treatment and a Bonferroni post-hoc test is displayed on each graph with the sexes grouped together. (*p < 0.05, **p< 0.01), n = 8-12 mice for each group. Results are expressed as the mean ± SEM.

### Microglial activation is specific to females following dsHMGB1 infusion to mPFC

The mPFC is often described as a key node for stress responsivity and emotional processing; these are key processes in the context of anxiety. Sex differences in stress-associated microglial activation are well documented (Bollinger et al., 2016). We sought to determine if microglia activation would be a shared phenotype between males and females underlying the similar behavioral outcomes. In a smaller cohort of mice, we repeated the implantation and dosing scheme as outlined in the previous section. 16 hours following the final infusion of dsHMGB1, mice completed the elevated plus maze assay and were immediately sacrificed for tissue collection. Again, an anxiety phenotype in the EPM was observed in males and females **(Suppl. Fig 3A and 3B)**. We triple stained Iba1, Tmem119, and CD68 to distinguish between surveillant and homeostatic, and reactive/phagocytic microglia respectively. We quantified the volume of Iba1 and Tmem119 **(Fig 2B, 2F, Suppl. Fig3C and 3D)**. There was no significant difference in Tmem119 volume alone for males or females; however, Iba1 volume differed significantly in females (p<0.05) **(Fig 2F)**. We then calculated the ratio of Tmem119 normalized to Iba1 for each reconstructed cell. We interpret this measurement as a ratio of homeostasis, as Tmem119 is often downregulated in the context of inflammation, injury, or disease progression (Spiteri et al., 2021). We observed a clear linear relationship between Iba1 volume and Tmem119 in control mice (**Suppl. Fig3E**). However, in dsHMGB1 treated mice we found this relationship was significantly altered in females but not males (**Fig 2C and 2G**). Female mice treated with dsHMGB1 showed decrease Tmem119:Iba1 relative to controls (p<0.05) **(Fig 2I)**. Next, we calculated phagocytic capacity by normalizing CD68 volume to Tmem119 and Iba1 merged volumes. Females but not males treated with dsHMGB1 had significantly increased phagocytic capacity (p<0.05) **(Fig 2J)**. Finally, we examined changes in morphology by calculating the number of Sholl intersections from concentric distances from the radius of each cell soma. In both males and females, dsHMGB1 decreases the number of intersections and we interpret this to reflect a decrease in complexity and an increase in reactivity **(Fig 2E and Fig 2I)**.

**Fig. 2.**
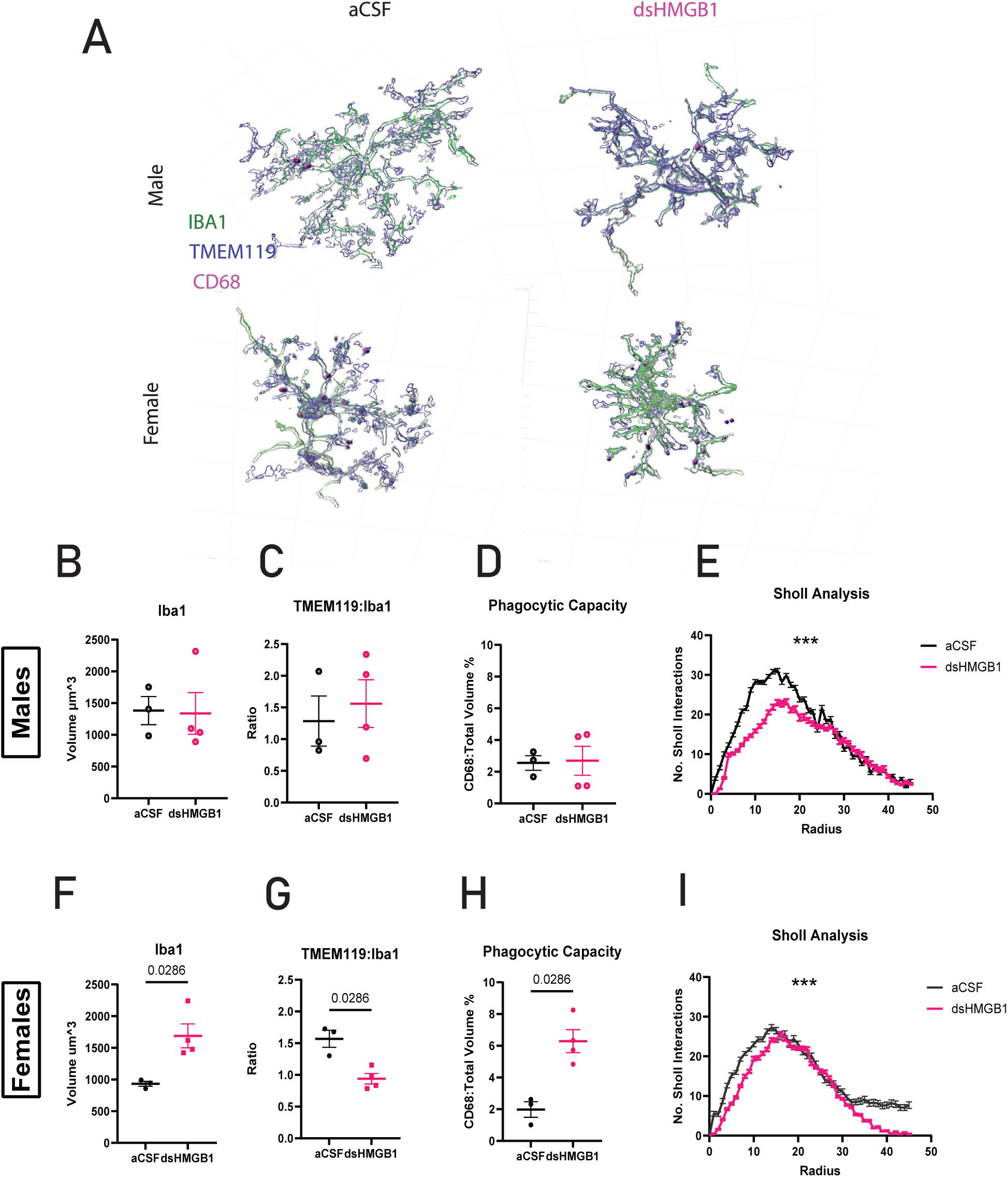
HMGB1 infusion induces morphological changes in female microglia. *A) Representative Images for IHC analysis (B-E) Microglial imaging analysis for male mice (B)Iba1 Volume Quantification (C) Tmem119:Iba1 Volume Ratio (D) Phagocytic Capacity Phagocytic capacity was calculated by normalizing total CD68 volume to the total volume of IBA1 and Tmem119 merged. (E) Male Microglia Have Significantly Altered Morphology following HMGB1 (F-I) Microglial imaging analysis for female mice. (F)Iba1 Volume Quantification (G) Tmem119:Iba1 Volume Ratio (H) Phagocytic Capacity Phagocytic capacity was calculated by normalizing total CD68 volume to the total volume of IBA1 and Tmem119 merged. (I)Female Microglia Have Significantly Altered Morphology following HMGB1 Statistical analyses were performed using Mann-Whitney* test *with the exception of Sholl Analysis. Statistical analysis for this measure was performed by comparing the Pearson r values of the aCSF condition and dsHMGB1 condition (\**p *< 0.05, \*\**p *< 0.01, n = 3-4 mice for each group, each mouse has an n=6-8 cells that were 3D reconstructed for analysis). Each dot represents an individual mouse. Results are expressed as the mean ± SEM.*

### dsHMGB1 exerts sex specific changes in RAGE/MyD88 signaling

Noting the sex-specific effects of dsHMGB1 on microglial measures, we hypothesized there could also be sex-specific changes in proinflammatory signaling pathways. HMGB1 is a putative ligand for TLR4 and RAGE receptors (Xu et al., 2019; Pluma et al., 2020). Canonical activation of either of these receptors requires subsequent activation of the adaptor protein MyD88. We utilized a CD11b+ cell specific isolation protocol where CD11b marks myeloid cells, which are solely expressed by microglia and macrophages in the brain. CD11b- and CD11b+ cell fractions were isolated from mice 16 hours after the final dose to maintain the same timeframe as behavioral and immunohistochemical experiments. All cells were isolated from the frontal cortex. In males, no significant changes were found in TLR4, RAGE, or MyD88. Female CD11b+ cells showed significant increases in RAGE **(Fig 3D)** and MyD88 **(Fig 3F)**. Surprisingly an increase in MyD88 was also observed in the CD11b-cells for dsHMGB1 treated females **(Fig 3F).**

**Fig. 3.**
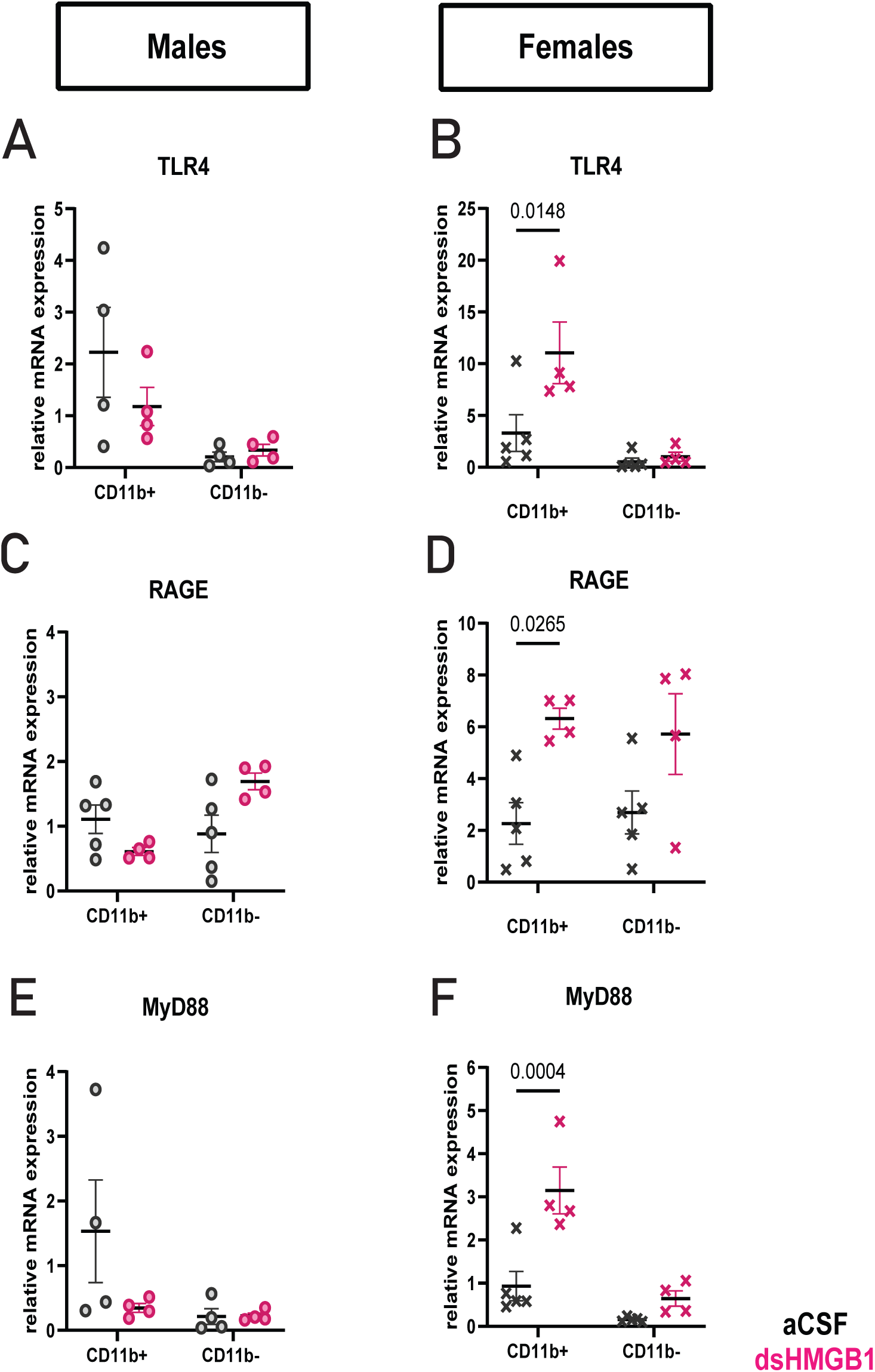
Male and Female Display Differential Transcriptional Responses to HMGB1. 16 hours following the final infusion of HMGB1, mice were sacrificed, and the frontal cortex was collected for cell isolation protocol as described in methods and materials section. CD11b+ cells are microglia and CD11b-cells are inclusive of neurons, astrocytes, and other glia (A) Male Expression of TLR4 (B) Female Expression of TLR4 (C) Male Expression of RAGE (D) Female Expression of RAGE (E) Male Expression of MyD88 (F) Female of Expression of MyD88. Statistical analyses were performed using Kruskal-Wallis Test (*p < 0.05, **p < 0.01, n = 4-5 mice for each group, each). Each dot represents an individual mouse. Results are expressed as the mean ± SEM.

### Microglial MyD88 is necessary for HMGB1 induced anxiety in females

To test the hypothesis that microglial MyD88 is necessary for anxiety-like behavioral responses to HMGB1, we bred our previously characterized mouse line that has conditional deletion of the MyD88 gene for CX3CR1 expressing cell types (Rivera et al., 2019), which includes all microglia in the CNS. The experimental workflow is the same as previous behavioral aims (Fig 1). Both male and female control mice responded with an anxiety-like phenotype following dsHMGB1 treatment as we have seen previously. cKO males looked similar to control mice, exhibiting anxiety-like behavioral responses after dsHMGB1 treatment. However, female cKO mice do not demonstrate an anxiety-like phenotype in the EPM or Novelty Suppressed Feeding (NSF) assay (**p < .01, 2-way ANOVA) **(Fig 4C and 4E)**.

**Fig. 4.**
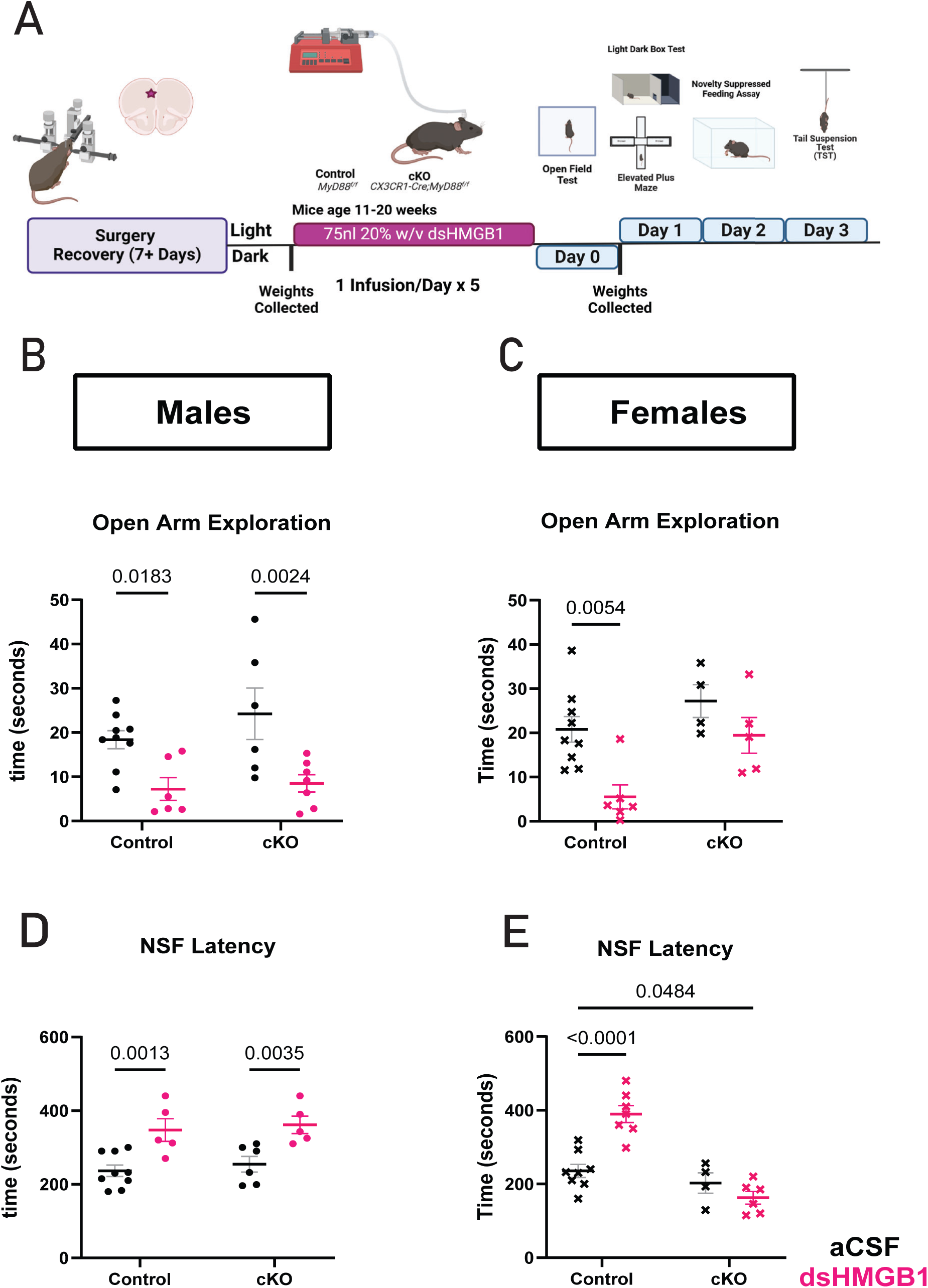
Microglial MyD88 is necessary for HMGB1 response in females. A) Schematic of Experimental Outline (Created with Biorender) B) Male Open Arm Exploration C) Female Open Arm Exploration D) Male Latency to feed in Novelty Suppressed Feeding Assay E) Female Latency to feed in Novelty Suppressed Feeding Assay Statistical analyses were performed using 2-way ANOVA and a Dunnett’s post-hoc values are displayed on each graph. (*p < 0.05, **p< 0.01), n = 5-10 mice for each group. Results are expressed as the mean ± SEM.

## Discussion

The hypothesis that inflammation is a critical mechanism which drives behavioral changes after stress exposure has substantial precedent in the literature (Won and Kim. 2020). Previous studies demonstrate that anti-inflammatory treatments can prevent and/or reverse behavioral deficits caused by stress in preclinical models (Basset et al., 2021). However, non-specific anti-inflammatory treatment is likely not sufficient or ideal to treat anxiety in humans (Hellmann-Regen et al., 2022). Therefore, more work is needed to unravel mechanisms of stress induced neuroinflammation to determine if therapeutics can be more precisely targeted.

HMGB1 preclinical models are emerging as a robust and reliable way to induce stress-associated behavioral effects (Huang et al., 2023). HMGB1-induced behavioral changes are reversed by antidepressants as well as the anti-inflammatory drug minocycline (Lian et al., 2017; Wang et al., 2019). More recent studies, including this one, demonstrate HMGB1’s utility as a pharmacological stressor which can impact specific behavior based on site of drug delivery. Moreover, HMGB1 does so while enacting other physiological changes associated with stress, e.g. weight loss in females, potentially providing novel insights into the pathophysiology of anxiety disorders.

Though anxiety and depression are twice as common in women, the current body of work related to stress, anxiety, and depression has been done primarily in male subjects (Kalin., 2020; Farhane-Medina et al., 2019). Pre-clinical stress models do not utilize female subjects to a comparable degree as male subjects nor to a degree that mimics the increased prevalence of these disorders in women (Lopez and Bagot, 2021). Various models of stress are utilized throughout the field, each exerting both specific and generalizing effects (Saxena et al., 2021). A now growing body of work is revisiting these canonical mechanisms of stress response in female subjects and uncovering both convergence and distinct mechanisms compared to males. In the present study our findings echo this trend; male and female adult mice infused with HMGB1 in the mPFC exhibit a shared anxiety-like phenotype in the EPM and NSF assay. However, the mechanisms were distinct, as changes in microglial reactivity were not evenly observed in both sexes.

Increased microglial reactivity may be a necessary mechanism by which stress alters behavior (Lehmann et al., 2019). Very little is established in the literature concerning understanding stress-associated HMGB1 mechanisms in a cell-specific manner. We hypothesized that microglial activation via PRRs would prove to be a crucial mechanism by which HMGB1 induces behavioral changes. We examined several previously identified hallmarks of stress-associated microglial pathology. Only females demonstrated significant alterations in microglia volume, phagocytic capacity, and homeostatic marker expression. Similarly, females specifically displayed significant increases in transcription of RAGE and MyD88 in microglial cells isolated from the frontal cortex. The role of microglial RAGE in the context of stress was first identified in male rats (Franklin et al., 2018). Here we demonstrate that adult female mice display increased activation of both microglial RAGE and MyD88 following dsHMGB1 infusion. Finally, we demonstrate that microglial MyD88 is necessary for HMGB1-induced anxiety only in female mice.

We must acknowledge several limitations with our study. Firstly, we infused 75nL of 20% dsHMGB1 into the mPFC once daily for five consecutive days. This dose was validated by its ability to significantly alter anxiety-like behaviors in males. We then utilized this same dosing scheme in the female experimental group. A previous study utilizing dsHMGB1 at a significantly higher dose and over a longer period reports a similar behavioral effect (Du et al., 2022). It is possible that some findings may be dose-dependent. Specifically, the transcriptional responses in male microglia trend in opposite directions than females. This was not surprising as it has been previously demonstrated that sub-chronic stress transiently downregulates RAGE and TLR4 (Franklin et al., 2018). However, it highlights the importance of potentially utilizing this model in both sexes across a broader dosing range to better delineate the tunability of HMGB1 responses. In addition, it is exciting to report results from the first utilization of dsHMGB1 as a pharmacological stressor in normally cycling females. Mice typically cycle across a period of 4-5 days. Both dosing and the initial behavior battery each occurred over a period of 4-5 days meaning mice were dosed and assayed at each point in their cycle. We were under-powered to investigate potential hormonal mechanisms more thoroughly in this study, but it is a possibility that warrants future investigation.

Finally, and possibly most importantly it would be exciting to draw broader conclusions about the role of microglial MyD88 in the physiological stress response, e.g. in response to psychological distress models. However, more work is needed to understand the role of microglial MyD88 in the stress response in such a way that generalizes across stress models.

There are several interesting future directions based on the findings of this study. Now that we have elucidated the importance of microglial MyD88 in female HMGB1 response, understanding the time course of resolution as well as investigating how this mechanism relates to stress-induced priming would be an interesting investigational aim. In line with this, we observed an almost bivalent effect of dsHMGB1 at this dose in both males and females. As previously stated, we utilized a significantly lower dose of dsHMGB1 that we believe allowed us to potentially capture a pseudo resilient v. susceptible phenotype. It would be interesting to follow up this observation with studies that examine how HMGB1-induced responses correlate with the response to stressors and subthreshold stressors.

This study adds to a growing body of work that demonstrates that similar behavioral response following stress may be the result of distinct sex-specific mechanisms. We also report the first use of HMGB1 as a model of anxiety in female adult mice. Moreover, our findings provide further evidence for the hypothesis that the ability of microglia to confer vulnerability to disease shifts from being male-biased in early life, to female-biased as we age (Lynch., 2022). Models of localized HMGB1 drug delivery could be utilized to further probe the effects of neuroinflammation with increased specificity. Expanding our understanding of mechanisms upstream of HMGB1 release as well as downstream signaling could continue to provide novel insights into stress pathology with potential broader implications. In conclusion, the data found in this study supports utilization of a local HMGB1 model to provide insight into pathological stress with great potential for future work and possible utility for therapeutic development.

## Materials and Methods

### Mice

MyD88-floxed mice were purchased from Jackson Laboratories (Bar Harbor, ME; Stock # 00888). Cx3Cr1-CreBT (MW126GSat) mice were generated and provided by L. Kus (GENSAT BAC Transgenic Project, Rockefeller University, NY). All methods for breeding and genotyping transgenic mice were conducted according to Rivera et al., 2019. Cx3Cr1-CreBT mice were crossed with MyD88-floxed mice over 3 generations until all offspring were fully MyD88-floxed (F/F) and either Cre negative (Cre 0/0: no modification to microglial MyD88) or Cre positive (Cre tg/0: removal of microglial MyD88), and then back-crossed onto a fully Jackson background. Genotyping of transgenic animals was conducted using polymerase chain reaction (PCR) on tail-snip DNA. Primer sequences used for PCR can be found in Table 1. Wild-type (WT) C57Bl/6J mice (used as stimuli for behavioral experiments) were purchased from Jackson Laboratories (Bar Harbor, ME; Stock # 000664). All animals were group housed in standard mouse cages under standard laboratory conditions (12-hour light/dark cycle, 23°C, 60% humidity) with same-sex littermates with free access to food and water.

**Table 1:**
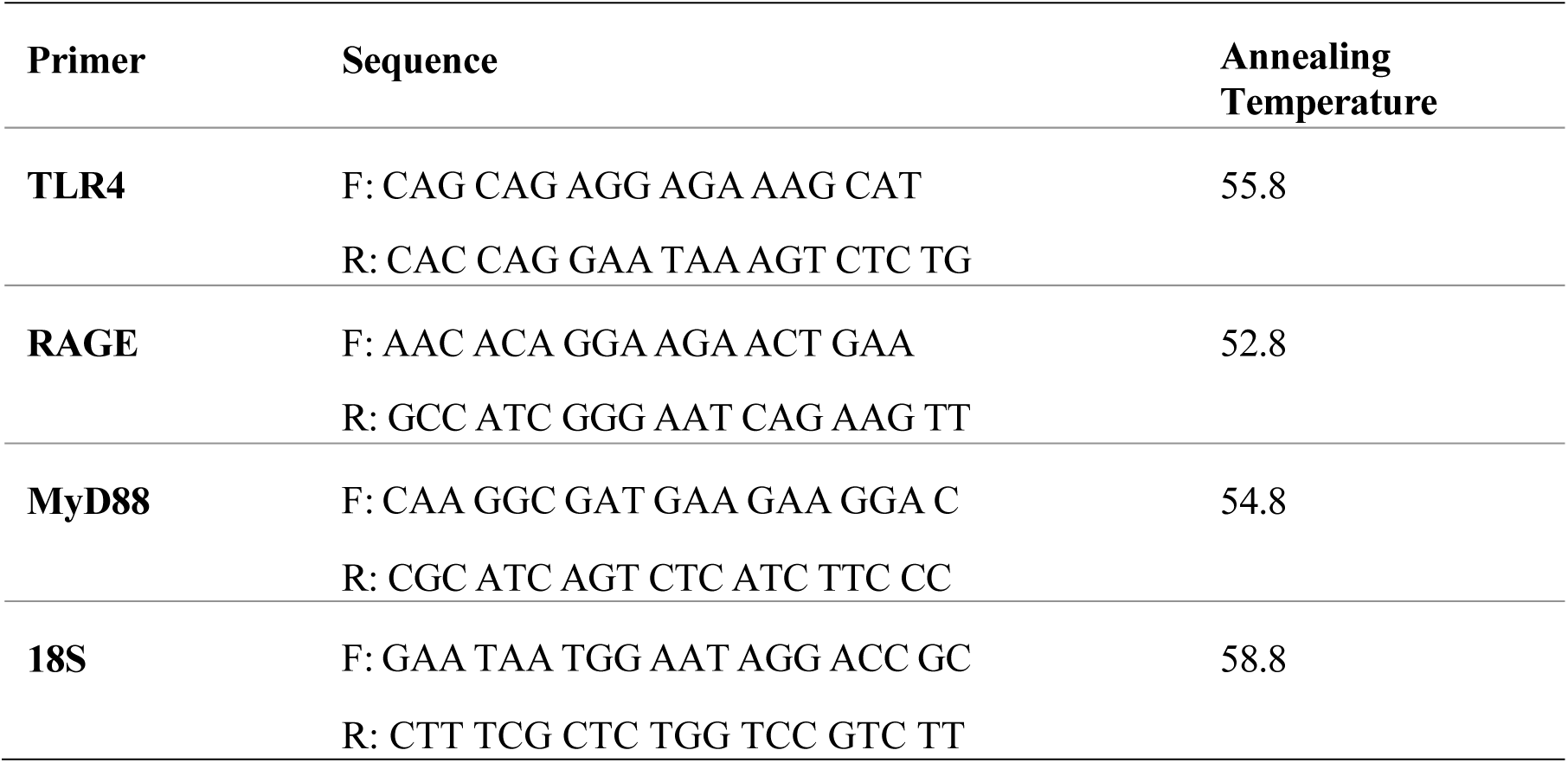
Primer Sequences for qPCR Experiments.

### Surgery

Mice were anesthetized with 1.5% isoflurane at a flow rate of 0.4 liter/min and stereotaxically implanted with an intracerebral ventricular (ICV) guide cannula (26G, 2.25mm from the pedestal; Plastics One, VA) into the left medial prefrontal cortex using the following coordinates from bregma (33): 1.5mm anterior-posterior, −0.4mm medial-lateral, and −2.25mm dorsal-ventral from the skull. After 10 days of recovery, the free moving mice were infused with 75nl of aCSF or recombinant HMGB1 (dsHMGB1 certified LPS free (0.2ug/ul); HMGBiotech) using a 26G internal cannula (Plastics One, VA) once daily for five consecutive days.

### Estrus Cycle Analysis

Smears were obtained by vaginal cytology collected at the end of behavioral tests and stained using the heamatoxylin-eosin method. Three cell types (nucleated epithelial cells, cornified epithelial cells and leukocytes) were counted to define the reproductive cycle (estrous), which is defined by the prevalence of each cell type: proestrus (nucleated), estrus (cornified), metestrus (all types in same proportion) and diestrus (leukocytes) (Westwood, 2008). Images were acquired with a light microscope Leica DM 4000 B (Leica, Wetzlar, Germany) with a ×10 objective lens (Plan 10×/0.25PH1).

### Weight Collections

Animals were weighed on the day of and the day preceding dosing. This weight was averaged to establish a baseline. Animals were weighed daily on dosing days and behavior days. The weight change reported for **(Fig 1E)** is the difference between baseline weight and weight 24 hours after dosing ends.

### Behavioral Tests

All behavioral tests were conducted approximately 12 hours apart from one another alternating between light and dark phase. Mice were habituated in the testing room for at least 30 min before the test. Examiners were blinded to the groups of mice. The testing room was sound-proof, and the temperature was maintained at 24–26°C. After each trial, the mice were put back into their home cages and the experimental apparatus was cleaned with 75% alcohol and Rescue cleaning solution to eliminate the odor that may affect animal behavior. Behaviors were tested in order consistent with experimental outlines provided in **Figure 1A and Figure 4A**.

### Open Field Test (OFT)

The OFT was performed in an open rectangular box with a bottom edge length of 50 cm and a height of 60 cm. The box was placed directly on the floor. After placing the mouse in the intermediate region of the box, animal behavior was recorded by a camera for 10 minutes. The bottom of the box was divided into the central zone (25× 25 cm^2^) and the rest of the outer zone by EthovisionXT. The time spent by the animal in each region as well as the moving distance in the box were calculated.

### Elevated Plus Maze (EPM)

The apparatus used for the EPM test was a cross-shaped device consisting of an intermediate platform region (5× 0.5 cm^2^) and two pairs of open arms and closed arms (25 × 5 cm^2^) connected to one another. The closed arms are surrounded by walls (16 cm), whereas the open arms have slight walls (0.5 cm). The entire maze was elevated to a height of 50 cm. The mouse was first placed in the intermediate region at the same position with its head toward closed arms, and its behavior in the device was then recorded by a camera for 5 min and the time spent by the animal in each arm was analyzed by EthovisionXT.

### Social Preference Assay

Social preference was measured using a two-chamber assay in which animals explored a novel object or a novel mouse as described in (Mague et al., 2022). The assay was run in a rectangular arena (61cm × 42.5cm × 22cm, L×W×H) fabricated from clear plexiglass with a clear wall separating the arena into two equal chambers with an opening in the middle allowing free access between both chambers. Plastic, circular holding cages (8.3cm diameter and 12cm tall) were centered in each of the two chambers and were used to hold either a novel object or sex- and age-matched C3H target mouse. The arena was evenly lit with indirect white light (∼125 lux). Plastic toys and glass objects were used as novel objects with the object being between 3– 5cm in all directions. For the test mouse, time spent interacting with a novel object and time spent interacting with the C3H mouse were measured. Social preference ratio was calculated by normalizing the social interaction time to the object interaction time.

### Novelty Suppressed Feeding

Following the completion of EPM, mice were food-deprived 18 hours prior to the test, with free access to water. On the day of testing, mice were moved to the dimly lit testing room one hour before the test. Mice were placed into one corner of a clear plexiglass open field apparatus (17 in. × 17 in. × 12 in.) with an opaque white acrylic floor covered with a thin layer of home cage bedding material. The room was maintained with red light conditions. A food pellet was placed in the center of the open field and mice were placed in one corner. Latencies to approach and to begin eating were recorded with a limit of 8 minutes. As soon as the mouse was observed to eat, or the 8-minute time limit was reached, the mouse was removed from the open field. And placed in the home cage and observed until it began to eat in the home cage.

### Home Cage Feeding Assay

Immediately following completion of the NSF trial. Mice were then placed in a cage with bedding material from their home cage to increase familiarity within the arena. Again, a mouse began the trial in the corner of the cage and the food pellet was placed in the center. Each trial was limited to 3 minutes and at the end of that time or after the mouse began its first feeding bout, the mouse was removed, and the trial completed. The latency to feed was recorded as the trial occurred and confirmed via video hand scoring.

### Light Dark Box Test (LDB) test

The apparatus for LDB test was a rectangular box comprising two connected chambers: a light open chamber with white walls (∼ 100 Lux, 15 × 20 × 25 cm^3^) and a dark covered chamber with black walls (∼ 5 Lux, 30 × 20× 25 cm^3^). The apparatus was placed on a table two feet off the ground. For every trial, the mouse was placed at the same position in the dark box. Animal behavior in the apparatus was then recorded for 8 min by a digital camera and the time the animal spent in each box was analyzed by EthovisionXT.

### Tail Suspension Test

For TST, mice were suspended from the side of a behavioral arena with white background for video tracking. At the beginning of the test, the mouse was fixed to the side of the arena with tape 1.5cm wide and approximately 15cm long. Mice were taped at the distal end of the tail and suspended 15 cm from the base surface. The camera was then turned on to record the animal behavior for 6 min. Immobility was defined by mobility (> 2.8%) and velocity parameters ( > 3 cm/s) set in EthovisionXT.

### Immunohistochemistry

Mice were perfused intracardially with ice-cold saline followed by phosphate-buffered 4% paraformaldehyde (pH 7.4) after anesthetized with CO2. The brain was removed and post-fixed overnight in the same fixative and dehydrated in 30% sucrose for 48 h at 4 °C. Coronal brain sections were cut at 30 μm by a cryostat (NX50, Thermo, USA). For immunohistochemical staining, the sections containing the mPFC, conventionally implicated in anxiety, were incubated in 0.1% Triton X-100 for 15 min and in 5% normal goat serum for 2 h firstly, and then were incubated with rabbit, chicken, or guinea pig anti-mouse primary antibodies against Iba1 (1:500, Abcam, UK) /TMEM119 (1:500, Millipore, USA) /Iba1 (1:500, Wako, Japan) or CD68 (1:500) overnight at 4 °C, then with anti-rabbit IgG-Alexa Fluor 488 or anti-rat IgG-Alexa Fluor 594 (1:1000, Invitrogen, USA) for 2 h at room temperature (RT). After repeated washing, the sections were then covered with glass coverslips and fluorescent images were captured by a confocal microscope (Olympus FV3000 inverted confocal). The analysis of fluorescence intensity and cell counting were performed by Image J software (NIH, USA). Microglia reconstructions and individual cell analysis was performed utilizing IMARIS 10.0 software. Structural analysis included the total length and the 3D arrangement of microglial branching using the Sholl analysis (number of intersections with concentric circles positioned at radial intervals of 1 μm). For each animal, 6-8 microglia from mPFC were analyzed and microglia from the same animal were averaged. Observers were blinded to the experimental condition of each subject.

### RNA Extraction

Samples were homogenized in Trizol® (Thermo-fisher scientific, NY) and then vortexed for 10 min at 2000 rpm. After 15 min resting at room temperature (RT), chloroform was added (1:5 with Trizol) and samples were vortexed for 2 min at 2000 rpm. Samples were next allowed to sit at RT for 3 min and then centrifuged (15 min at 11,800 rpm; 4°C). This resulted in a gradient from which the top, clear, aqueous phase was separated and placed into a fresh tube. Isopropanol was then added to the new tube to precipitate RNA (1:1 with aqueous phase) and samples were again vortexed, allowed to set at RT for 10 min, and then centrifuged (15 min at 11,800 rpm; 4°C). Pellets obtained after this step were rinsed twice in ice-cold 75% Ethanol and then resuspended in 8 μl of nuclease-free water. RNA was frozen at −80°C until cDNA synthesis and qPCR. A NanoDrop Spectrophotometer (Thermo Scientific, Wilmington, DE) was used to determine RNA quantity and purity. RNA was considered pure enough for further use based on 260/280 (RNA:protein; range: 1.8–2.0) and 260/230 (RNA: Ethanol; range 1.6– 2.0) ratios.

### qPCR

cDNA was synthesized using the QuantiTect Reverse Transcription Kit (Quiagen, Hilden, Germany). Briefly, 200ng of RNA/12 μl of nuclease-free water was pre-treated with gDNase at 42°C for 2 min to remove genomic DNA contamination. Next, master mix containing both primer-mix and reverse transcriptase was added to each sample and all samples were heated to 42°C for 30 min and then 95°C for 3 min (to inactivate the reaction) in the thermocycler. Quantitative real-time PCR (qPCR) was conducted using QuantiTect SYBR Green PCR Kit (Quiagen, Catalog #:204057, Hilden, Germany. To examine HMGB1-dependent signaling that is mediated by the MyD88-dependent pathways, we selected 3 genes within the signaling cascade of HMGB1 including putative receptors RAGE, TLR4, as well as MyD88 (Magna & Pisetsky, 2014). qPCR primers were designed in the lab and ordered from Integrated DNA technologies (Coralville, IA).

PCR product was monitored using a Mastercycler ep *realplex* (Eppendorf, Hauppauge, NY). Male and female samples were run on separate plates, and thus, cannot be directly compared. I8S was used as a house-keeping gene and relative gene expression was calculated using the 2−ΔΔ^CT^ method, relative to 18S and the lowest sample on the plate (Williamson et al., 2011; Livak & Schmittgen, 2001).

### Statistical Analysis

All data are expressed as the mean ± SEM. The required sample sizes were estimated based on our experience and Power analysis was used to justify the sample size. Statistical analysis was conducted by GraphPad Prism10.2 (GraphPad Software, USA). The Shapiro–Wilk test was used to assess whether the data followed a normal distribution. If the data was normally distributed, one-tailed paired or unpaired Student’s *t* test was used for comparison between two groups; if not, Mann–Whitney test was used instead. For groups with a normal distribution, a two-way ANOVA was utilized to assess the significance of two factors and the interaction of those factors (ie: sex and treatment) either Bonferroni or Dunnett’s post-hoc test was used. The significance level was set at *P* < 0.05.

## Supporting information

Detailed Statistics

## Author Contributions

AR conceived the study and together with KD and SDB designed the experiments. AR, DN, JD, DA, and MC performed the experiments. AR analyzed all data and wrote the manuscript together with KD and SDB. KD and SDB oversaw the project.

## Acknowledgements

We would like to acknowledge Michael Patton, Ph.D. and Lauren Green Ph.D. for training in key techniques.

---

**Supplemental Figure 1.**
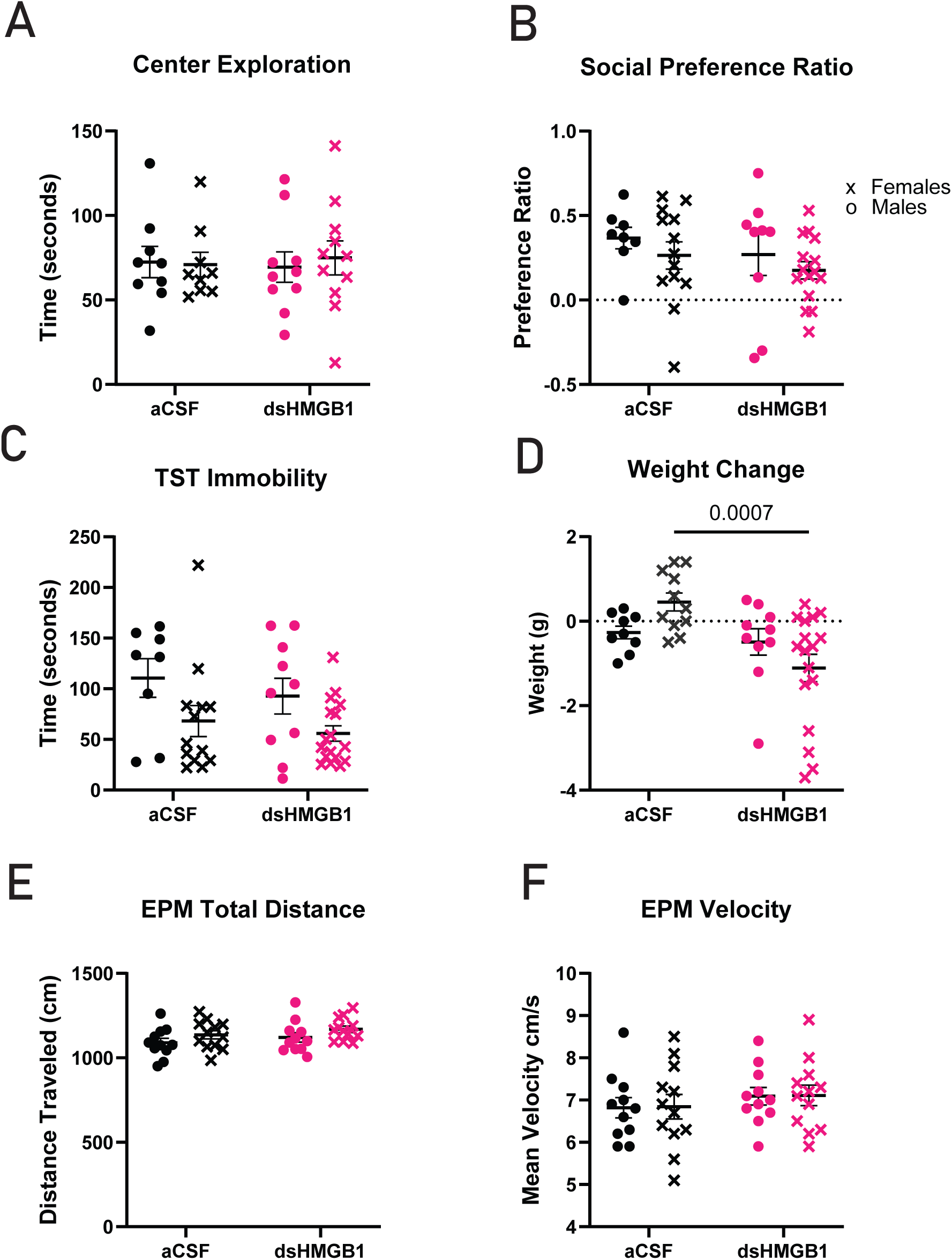
HMGB1 does not impact OFT, Social Preference, or TST immobility. A) Center Exploration of the OFT is unchanged by HMGB1 B) Social Preference is not altered by HMGB1 C) TST Immobility is not significantly effected by HMGB1 D) Weight Change in response to HMGB1 is sex-specific E) Total Distance Traveled in EPM F) Mean Velocity in EPM Statistical analyses were performed using 2-way ANOVA to confirm primary effect of treatment and a Bonferroni post-hoc test is displayed on each graph (*p < 0.05, **p < 0.01, n = 8-12 mice for each group Results are expressed as the mean ± SEM.

**Supplemental Figure 2.**
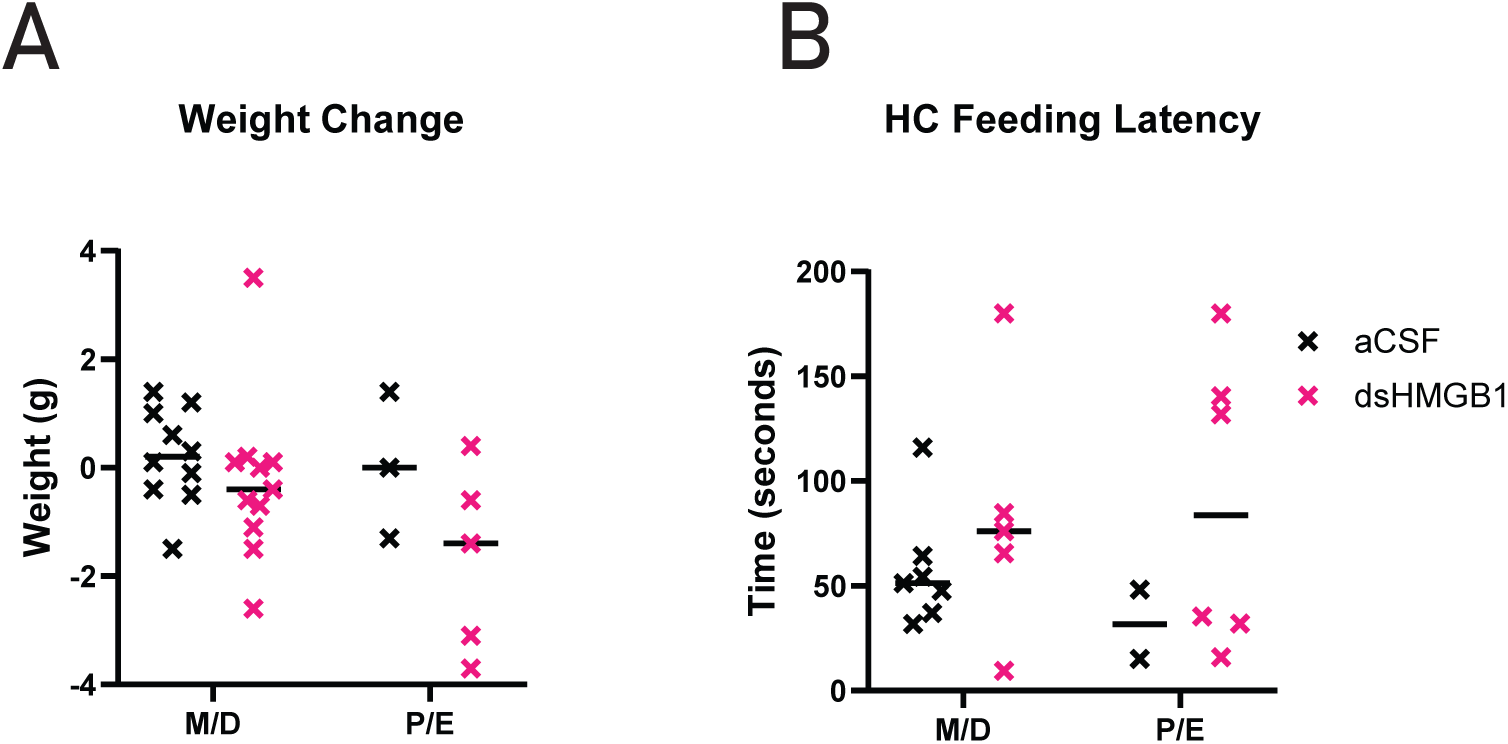
Relationship Between Estrus Cycle and Behavioral Effects of HMGB1. A) Weight Change and Estrus Cycle following HMGB1 B) Home Cage Feeding Latency and Estrus Cycle following HMGB1 Statistical analyses were performed using 2-way ANOVA test (*p < 0.05, **p < 0.01, n = 22). Results are expressed as the median for each group.

**Supplemental Figure 3.**
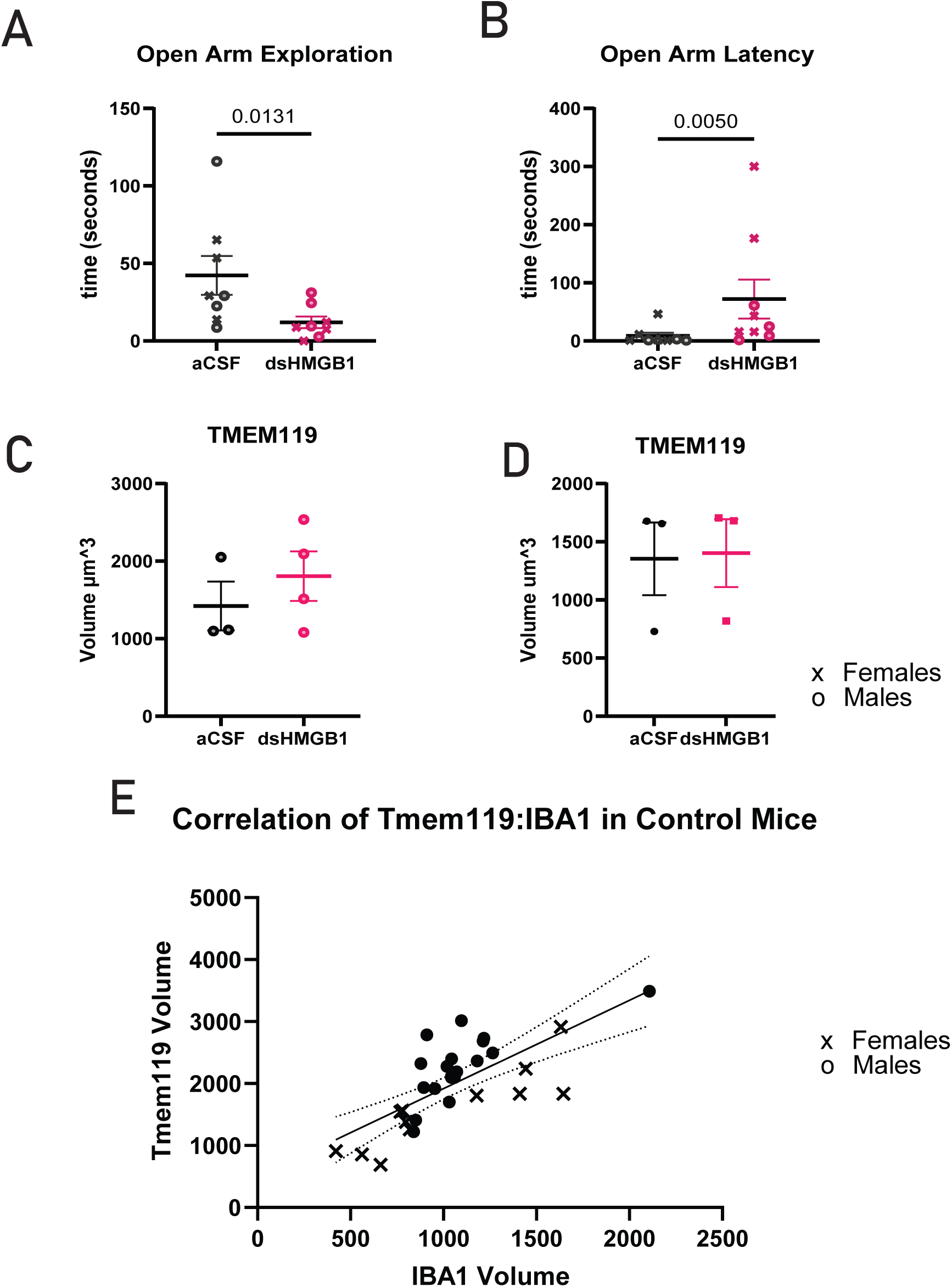
HMGB1 induced anxiety is robust in cohort utilized for microglia imaging experiments. (A) Open Arm Exploration for animals used for IHC imaging experiments (B) Latency to Explore Open Arms for animals used for IHC imaging experiments, n = 3-4 mice for each group. Each dot represents an individual mouse. Results are expressed as the mean ± SEM. (C) Male Tmem119 Volume Quantification (D) Female Tmem119 Volume Quantification (E) Linear Regression of TMEM119 volume and Iba1 volume for control mice. Individual cells, 5-6 per mouse are plotted for controls. The equation of the line is y= 1.424x+494.8. Pearson correlation coefficient, r= 0.727. p < 0.0001.

**Supplemental Figure 4.**
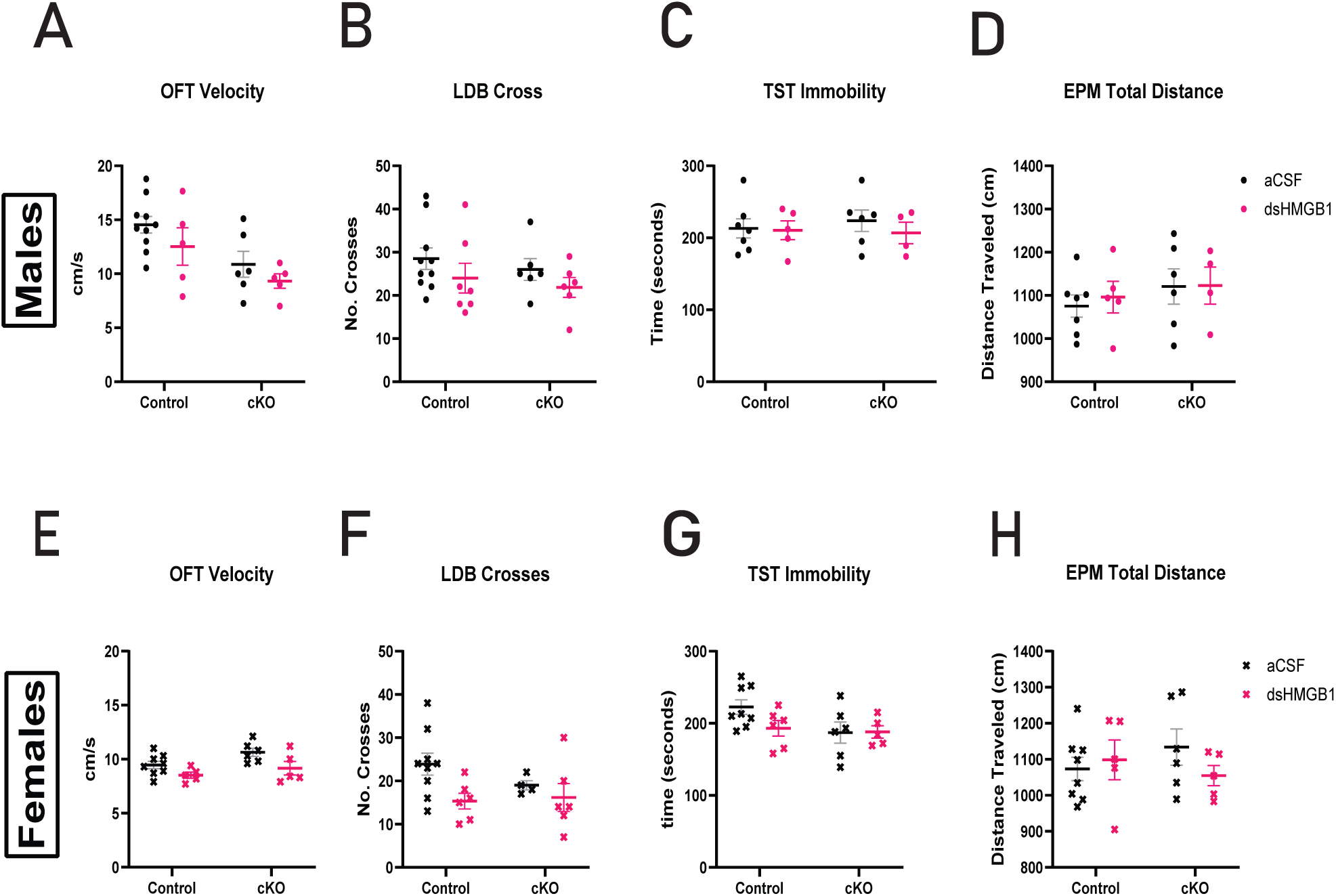
No Changes in OFT, LDB, TST, or Distance traveled in EPM were observed in response to dsHMGB1. A-D) Male Behavioral Data A) OFT Velocity is not changed by HMGB1 B) Light Dark Box (LDB) crossings are not impacted by HMGB1 C) TST Immobility remains constant following HMGB1 D) EPM Total Distance Traveled E-H) Female Behavioral Data E) OFT Velocity is not changed by HMGB1 F) Light Dark Box (LDB) crossings are not impacted by HMGB1 G) TST Immobility remains constant following HMGB1 H) Total Distance Traveled in EPM *p < 0.05,**p < 0.01 N per group is 5-10, per treatment group. Statistical analyses were performed using 2-way ANOVA.

